# Stalling chromophore maturation of the fluorescent protein Venus reveals the molecular basis of the final oxidation step

**DOI:** 10.1101/2020.10.13.337386

**Authors:** Husam Sabah Auhim, Bella L. Grigorenko, Tessa Harris, Igor V. Polyakov, Colin Berry, Gabriel dos Passos Gomes, Igor V. Alabugin, Pierre J. Rizkallah, Alexander V. Nemukhin, D. Dafydd Jones

## Abstract

Fluorescent proteins (FPs) have revolutionised the life sciences but the mechanism of chromophore maturation is still not fully understood. Incorporation of a photo-responsive non-canonical amino acid within the chromophore stalls maturation of Venus, a yellow FP, at an intermediate stage; the crystal structure reveals the presence of O_2_ located above a dehydrated enolate imidazolone (I) ring, close to the strictly conserved Gly67 that occupies a twisted conformation. His148 adopts an “open” conformation, potentially allowing O_2_ access to the chromophore. Absorption spectroscopy supported by QM/MM simulations suggest that the first oxidation step involves formation of a hydroperoxyl intermediate in conjunction with dehydrogenation of the methylene bridge. A fully conjugated mature chromophore is formed through release of H_2_O_2_ upon irradiation of this intermediate, both *in vitro* and *in vivo*. The possibility of interrupting and photochemically restarting chromophore maturation, and the mechanistic insights opens up new approaches for engineering optically controlled fluorescent proteins.

## Introduction

Fluorescent proteins (FPs) represent a unique family of proteins that emit light in the visible region of the spectrum without the requirement of any additional cofactor, ^1–4^ so enabling them to act as genetically encoded markers and sensors ^5,6^. Formation of the chromophore occurs through the stepwise covalent arrangement of three contiguous residues Xaa-Tyr-Gly (where Xaa can be various amino acid residues) in the presence of molecular O_2_ (Figure 1a). In the case of Venus, these three residues are 65-Gly-Tyr-Gly-67 (Figure 1a). Venus is a yellow fluorescent derivative of the classical *Aequorea victoria* GFP protein ^7^. Amongst the various mutations introduced to generate this yellow variant, two key changes are the introduction of glycine at residue 65 (from serine) and tyrosine at residue 203 (from threonine); Tyr203 pi-stacks with the chromophore causing a red shift in excitation and emission into yellow region ^7,8^. The mature chromophore has an extended conjugated bond network comprised of the aromatic P ring and the imidazolone I ring linked via a methylene bridge (see Figure 1a for ring and atom notation), which is located within the core of the β-barrel structure ^1,4,9,10^. The first step in chromophore formation is cyclisation through linkage of the backbone amine of the strictly conserved Gly67 residue to the carbonyl carbon of the Xaa 65 residue (Gly65 in the case of Venus). Cyclisation is followed by dehydration/O_2_-dependent oxidation steps to generate a fully conjugated system. There has been much discussion about the exact mechanism of chromophore formation, especially the order of the dehydration and dehydrogenation/oxidation steps ^4,9,10^, and the role O_2_ plays in the rate-limiting oxidation step ^11^. The availably of structures representing trapped intermediates has helped shed light on potential mechanisms ^12–16^ but there are still questions to be addresed. To our knowledge, neither the crucial O_2_ nor the oxygenated intermediates have been observed, while theoretical simulations ^17^ have speculated on the mechanism of action. Gly67 is strictly conserved and critical to maturation ^15,18^, but its role is unclear.

**Figure 1.**
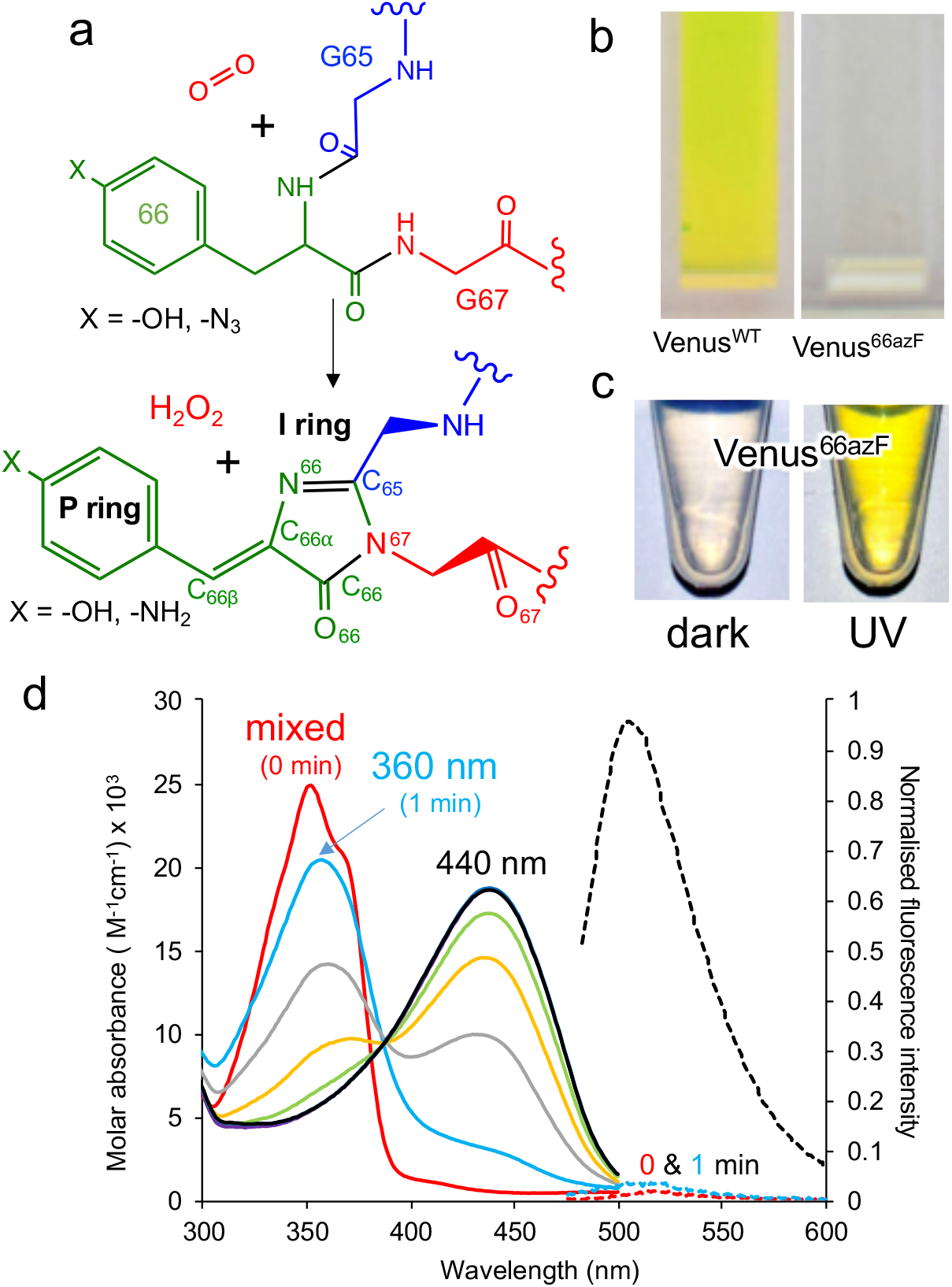
Effect of azF incorporation at residue 66 of Venus. (a) Scheme outlining the basic maturation of Venus. The bond and ring nomenclature are described in the lower structure. (b) Solution colours of Venus^WT^ and Venus^66azF^. (c) Solution colours of Venus^66azF^ before (dark) and after (UV) illumination with UV light. (d) Absorbance (solid line) and fluorescence (dashed line) of Venus^66azF^. Red, light blue, grey, orange, green, purple, dark blue represent spectra after 0, 1, 5, 10, 15, 30, 45 and 60 min. Full spectral properties are shown in Supporting Table S1. Full fluorescence emission time course and *in vivo* imaging are shown in Supporting Figure S1.

Manipulating the chemical properties of the chromophore by protein engineering, either directly through changes to two of the three chromophore residues, or indirectly through changing the chromophore environment, have generated a range of new fluorescent proteins, including Venus ^7^, with properties suited to their particular application ^6^. One of the most important FP class for super-resolution imaging is the photo-controllable FPs, whereby fluorescence is either switched on/off, or spectral properties significantly shifted in response to light ^19–22^. Mechanisms of action involve chemical modifications such as decarboxylation of Glu222 (e.g. PA-GFP ^23^), backbone cleavage (e.g. Kaede ^24^) and chromophore hydration (e.g. Dreiklang ^25^), or conformational changes such as chromophore *cis/trans* isomerisation (e.g. Dronpa ^26^, rsGFP ^27^). The use of photochemically active non-canonical amino acids (ncAA) has further expanded optical control approaches ^28^. Phenyl azide photochemistry is particularly useful as it has been used to turn on, off, or switch the fluorescence properties of green ^29,30^ and red ^31^ FP types.

Here we use the photochemical properties of genetically encoded phenyl azide to stall chromophore maturation of Venus at an immature non-fluorescent intermediate (termed im-Venus^66azF^) state before UV irradiation instigates maturation to a final fluorescent form. The structure of the intermediate reveals the protein has undergone the dehydration but not the oxidation step. Additional new structural features add further new insights, including an essential role for the strictly conserved Gly67 and, for the first time, experimental observation of molecular O_2_ in direct proximity to the immature chromophore. The combination of experimental spectroscopy with quantum mechanical/molecular mechanics (QM/MM) simulations allowed us to propose a mechanism for the O_2_ dependent oxidation step whereby a hydroperoxyl intermediate is formed as part of the oxidation mechanism.

## Results

### Incorporation of phenyl azide chemistry within the Venus chromophore

To incorporate a photochemical switch into Venus, we replaced the chromophore residue Tyr66 with the aromatic non-canonical amino acid *p*-azido-L-phenylalanine (azF) ^32,33^; effectively the hydroxyl group of tyrosine is replaced with an azide group. The variant termed Venus^66azF^ is produced as a colourless protein (Figure 1b) with no inherent fluorescence (Figure 1b-d and S1), and acts an excellent starting point for an optically controlled FP. The absorbance spectrum of the colourless non-fluorescent Venus^66azF^ reveals the protein absorbs in the UV region. Absorbance peaks at ~350 nm with a significant shoulder at ~360-370 nm; no fluorescence is observed from this species (Figure 1d and S1a-b). Irradiation of Venus^66azF^ with near-UV light converts the protein to a coloured and fluorescent form both *in vitro* (Figure 1c-d, S1a-b) and *in vivo* (Figure S1c-d), with an absorbance λ_max_ shifting to 440 nm (Supporting Table S1). Fluorescence emission increased over 100-fold on activation, allowing imaging of bacterial cells expressing the activated Venus^66azF^ (Figure S1d). Spectroscopic analysis of the time course of activation reveals a potential two-step process. Within 1 min of UV exposure, the double hump absorbance spectrum feature disappears, with the remainder of the time points forming a clear isosbestic point at ~390 nm suggesting conversion directly from one form to another. As the initial dark sample diverges away from the isosbestic point, it suggests a primary conversion to an intermediate species, which is also non-fluorescent, followed by a slower conversion to the activated form.

### Structure of im-Venus^66azF^

The structure of the pre-photoactivated Venus^66azF^ (from herein referred to as immature or im-Venus^66azF^) was determined (see Supporting Table S2 for structural statistics) to gain an insight into the molecular nature of the non-fluorescent, immature chromophore. The general structure of im-Venus^66azF^ is similar to mature Venus (henceforth termed Venus^WT^), with a root-mean-squared deviation (rmsd) over the backbone of 0.39 Å (Supporting Figure S2a). The most significant differences occur in and around the chromophore (termed CRO). Electron density fits well to a cyclised chromophore containing the intact azide group present on the P ring (Figure 2 and S2b). The lack of the electron density protruding from the I ring at the C_65_ position suggests the dehydration step in chromophore maturation has occurred by this point (Supporting Figure S2b). Compared to Venus^WT^, the I ring component shifts position in im-Venus^66azF^ due to the more acute angle in the methylene bridge (Figure S3b). A comparison of the residues surrounding the CRO shows that several residues exist in different conformations compared to Venus (Figure S3). His148, a critical H-bonding residue to the mature CRO, exists fully in the open-gate conformation together with Glu222, another residue necessary for function, and Tyr203 (pi-stacks with the CRO in Venus^WT^) also shift position with respect to the CRO (Figure 2a).

**Figure 2.**
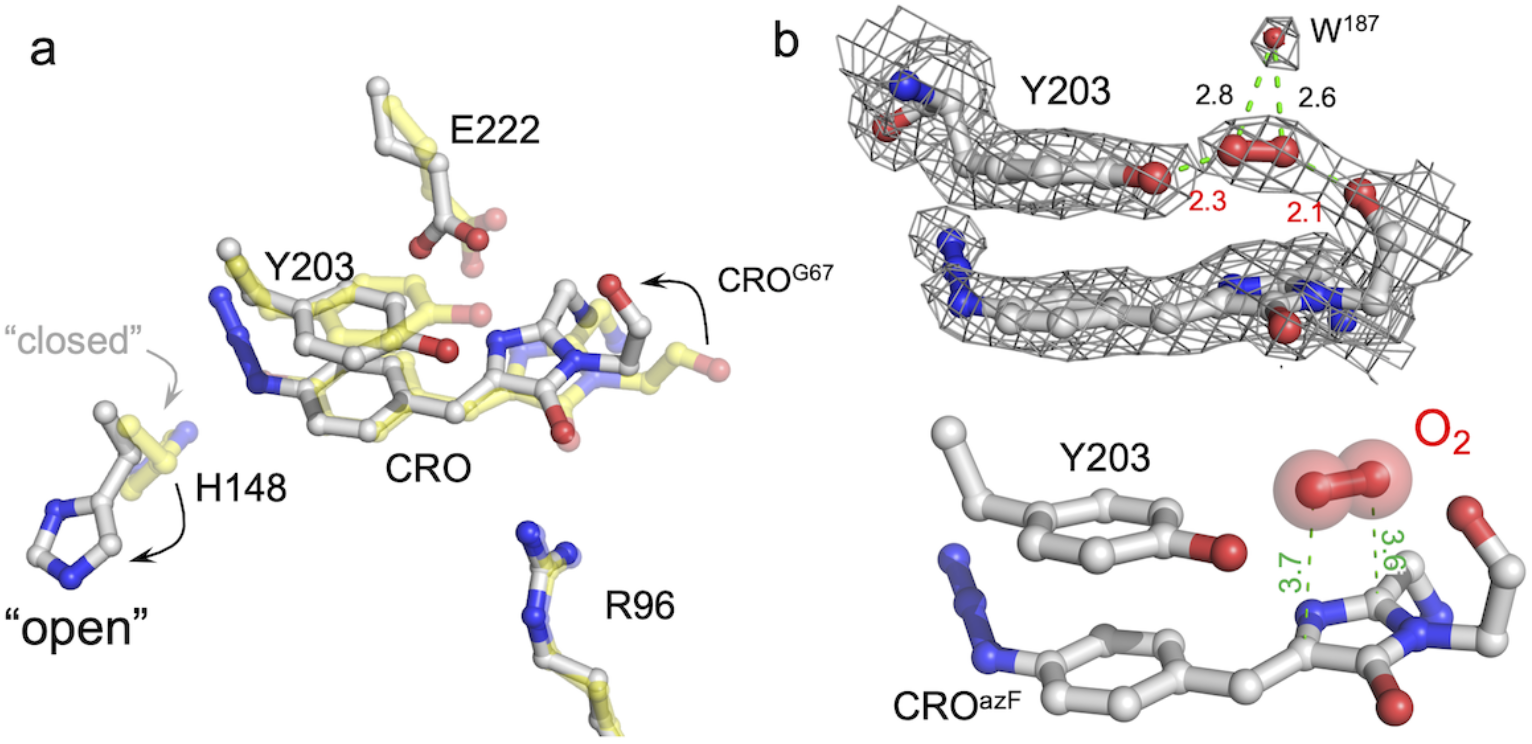
Structure of im-Venus^66azF^ proximal to the chromophore. (a) comparison of im-Venus^66azF^ (grey; PDB 6sm0) with Venus (yellow; PDB 1mwy ^8^). CRO is the chromophore (Gly65-Tyr/AzF66-Gly67). (b) Position of the O_2_ molecule in im-Venus^66azF^. The top panel shows the electron density (2Fo-Fc, 1.5 sigma) for the CRO, Y203 and O_2_ together with an additional water molecule. The lower panel removes the electron density for clarity. Relevant distances are shown in Å.

Further analysis of the chromophore reveals several novel features that provide us with insights into CRO maturation and the role of molecular O_2_ in the process. In im-Venus^66azF^, the backbone carbonyl of the strictly conserved Gly67 is ~180° out of position compared to that observed in other FPs (Figure 2a-b). The C-O bond is also longer (1.36 Å) than that generally observed for a carbonyl C-O bond within the context of a peptide bond (~1.24 Å). To our knowledge, the only other time this conformation has been observed is in the unpublished structure of an immature chromophore of a GFP maturation disabling mutant determined by the Getzoff group (PDB 2qt2; Supporting Figure S4). There is additional electron density sandwiched between the Gly67 carbonyl group and the hydroxyl group of Tyr203 (Figure 2b), which we have assigned to molecular O_2_ after attempting to: (1) refine the structure with two H_2_O but the presence of Tyr203 effectively blocks the ability of two water molecules to occupy this position; (2) refinement of im-Venus^66azF^ with one water molecule left an elongated tail of density; (3) molecular O_2_ fitted best. The O_2_ molecule lies between the carbonyl oxygen of Gly67 and the hydroxyl group of Tyr203 above the plane of I ring element (Figure 2b). O_2_ has been postulated to be positioned either above the plane of the chromophore facing Glu222/Tyr203 or below the chromophore plane facing Arg96; here it is clear O_2_ above the plane of the chromophore on the Glu222/Tyr203 face (Figure 2b).

Bond lengths and angles also provide an insight into the structure of the trapped chromophore intermediate (Figure 3; Supporting Table 3). During maturation, O_2_ is thought to be involved in generating the final C=C that links the P-ring and I ring (C_66β_-C_66α_ to C_66β_=C_66α_). In Venus, the C_66β_-C_66α_ bond is 1.35 Å, as expected for a C=C while the same bond is 1.45 Å im-Venus^66azF^, similar in length to adjacent C_66γ_-C_66β_ in both proteins (Figure 3a). This suggests that the C_66β_-C_66α_ in im-Venus^66azF^ remains as a single bond. The bond angle between C_66γ_-C_66β_-C_66α_ is also more acute for im-Venus^66azF^ (118° *versus* 131° for Venus; Figure 3). In the I-ring, the C_66_-O_66_ bond is 1.48 Å for im-Venus^66azF^, which is longer than would be expected for a keto-carbonyl C=O bond (1.20 Å) as observed in Venus (Figure 3b) and a standard C-O single bond (1.43 Å). This longer C-O bond also makes polar contacts with the critical maturation Arg96, a residue that occupies a near-identical position in Venus^WT^ (Figure 2a). Based on these measurements, we predict that the enolate is the most likely form of C_66_-O_66_ (Figure 3a), as has been proposed previously for the dehydrated intermediate ^16^, with Arg96 stabilising the negative charge. The associated C_66_-N_67_ bond is shorter (1.26 Å) compared to Venus (1.36 Å). The C_66_-N_67_ constrained within the cyclic configuration could indicate that this section of the ring is fixed as an ^-^O-C=N arrangement. Analysis of the rest of the bond lengths comprising the I ring (Figure 3a and Table S3), together with the trigonal planar arrangement of the I ring with the C_66β_, suggests that there may be a delocalised system around the I ring, with the positive charge spread around the ring.

**Figure 3.**
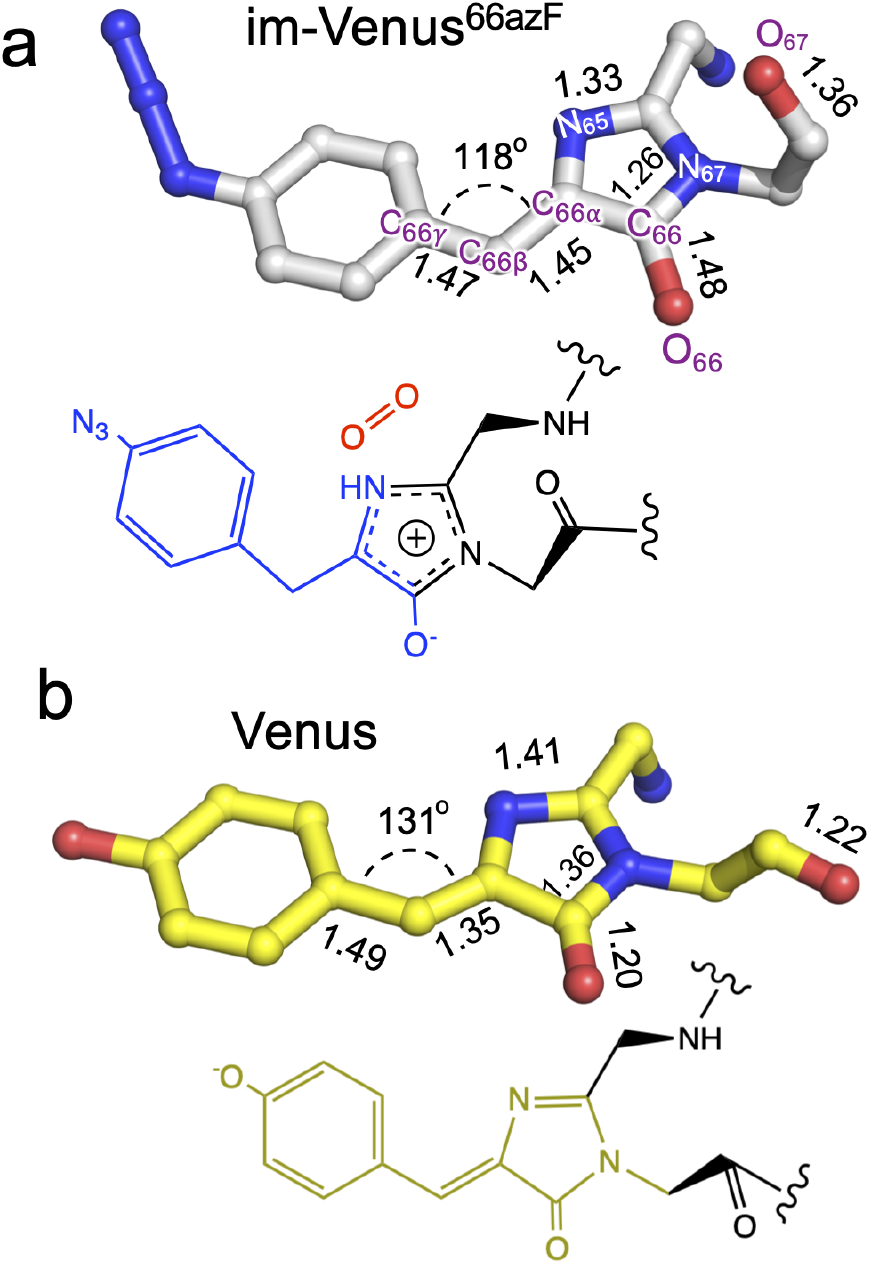
Chromophore bond distances and angles for im-Venus^66azF^ and mature Venus^WT^ fluorescent proteins. (a) im-Venus^66azF^ with proposed chemical structure; (b) Venus^WT^ (PDB code 1myw ^8^) with the chemical structure. A full list of bond lengths can be found in Supporting Table S3.

### Simulation of the oxidative step of chromophore maturation

To correlate the structural and functional observations, modelling of potential intermediates was performed using the atomic coordinates of im-Venus^66azF^ obtained in this work as a starting point and was associated with the observed absorbance data. The overall scheme based on the simulations is shown in Figure 4. The simulations predict that the first step is Gly67 switches to its energetically more favourable (~7 kcal/mol) canonical configuration. Triplet state oxygen can now access the I ring with concomitant protonation of Glu222, which acts as a general acid/base in the maturation scheme. The oxidation steps then proceed starting from partial negative charge transfer to O_2_, which switches from the triplet to singlet state. Glu222 is protonated with N_65_ donating the proton (Figure S5). The O_2_ then attacks C_65_ (and not C_66α_) generating a peroxy intermediate (Figures 4 and S5). The peroxy anion then abstracts a proton from C_66β_ to form the stable hydroperoxyl intermediate with Glu222 protonating N_65_ (Figure 4 and S5); the hydroperoxyl species has a predicted absorbance of 360 nm. As well as Gly67 converting through to its canonical conformation, the formation of the methylene bridge between the I and P rings results in a shift to a configuration similar to that observed for the mature chromophore observed in Venus^WT^; the methylene bridge bond angle is now 134° (Figure 5a).

**Figure 4.**
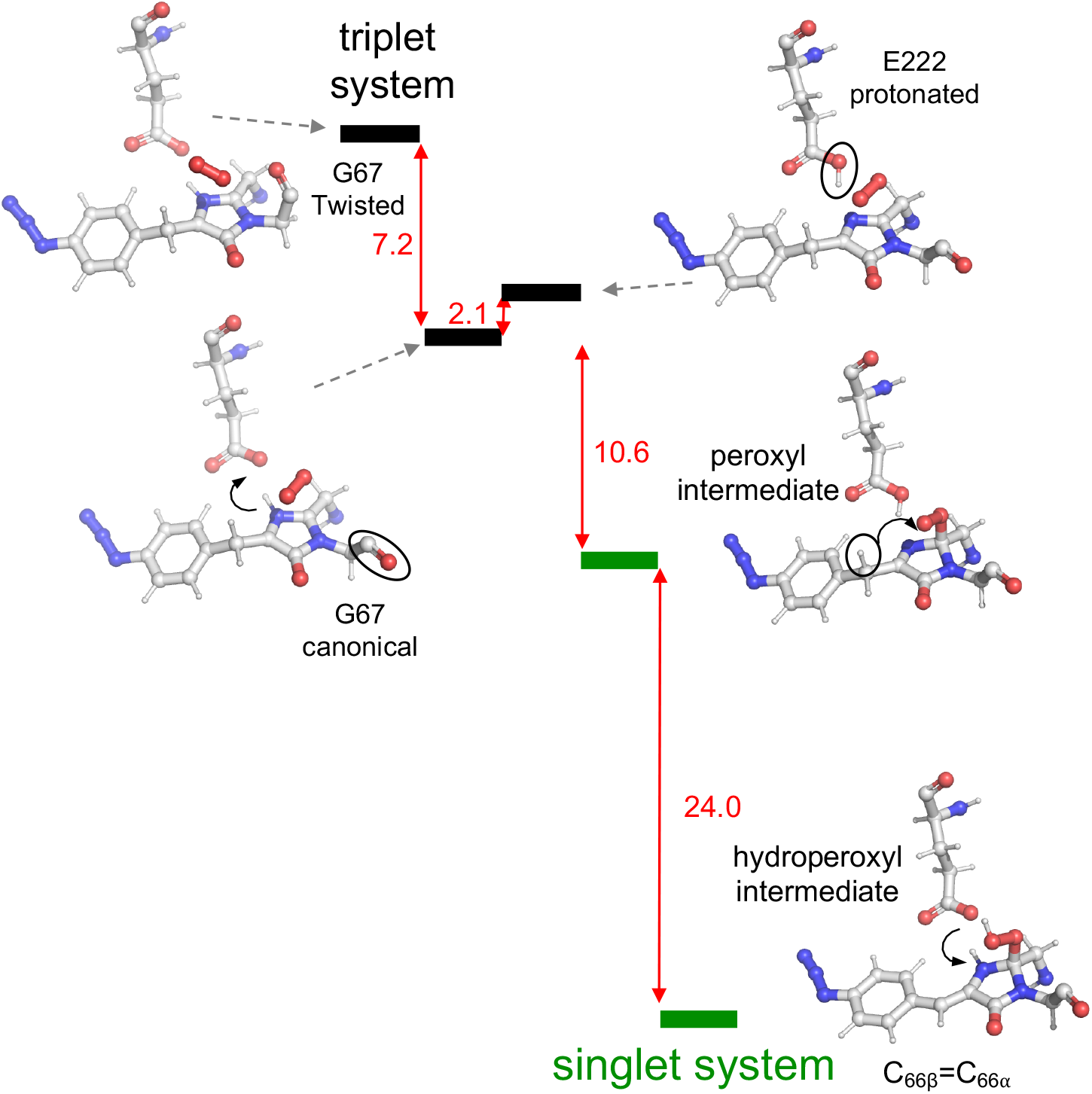
Reaction scheme for oxygenation of the chromophore as predicted by QM/MM modelling. Triplet and singlet systems are differentiated by black and green lines, as indicated in the diagram. The ΔE between each state is shown in red with units of kcal/mol. Significant changes between each state are outlined in the diagram.

**Figure 5.**
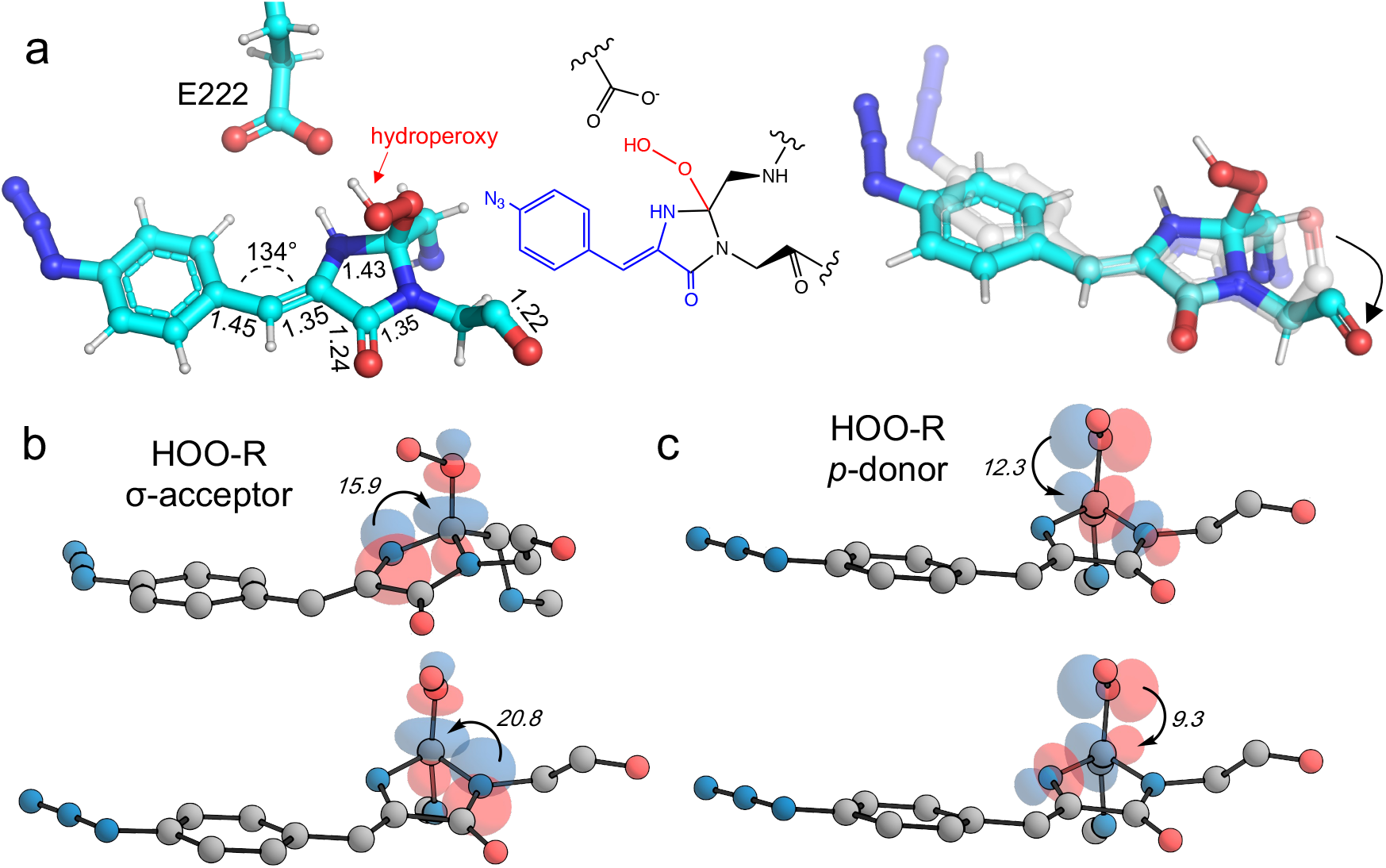
Models of the hydroperoxyl intermediate. (a) Model structure as predicted by QM/MM simulations with the hydroperoxy group bound to C_65_ as indicated. The chemical structure of the intermediate is also shown. On the right-hand side is a comparison of the modelled hydroperoxyl intermediate (cyan) with the crystal structure of im-Venus^66azF^. (b-c) Stereoelectronic interactions in hydroperoxyl intermediate (hydrogen atoms omitted for clarity with the numbers related to energies in kcal/mol). Both the hydroperoxyl moiety acting as a σ-acceptor (b) and *p-* donor (c) are shown.

The lone pair of the hydroperoxyl group (-OOH) in the model has near-ideal alignment with the two C-N bonds (Figure 5c-d). This favourable stereoelectronic arrangement activates stabilizing hyperconjugative n_O→σ*CN_ interactions, which can partially compensate for the loss of aromatic stabilisation in the I ring. This is complemented by two strong n_N→σ*CO_ interactions. The importance of the latter effect is expected to grow further in the transition state for the final C-O bond scission where it provides an important transition state stabilization effect that can significantly assist the final aromatizing step of the cascade. ^34–36^

The final step occurring over the irradiation cycle is the conversion to a fully mature fluorescent chromophore. Two events need to be considered: reduction of the azide and full conjugation of the chromophore through a loss of the hydroperoxyl moiety. The spectral properties suggest that the final end product is likely to be the phenylamine form of the mature chromophore as has been observed before (Table S1 and ^30^). Simulations concur with this with the final product having a predict absorbance max at 444 nm, close to the 440 nm observed in Figure 1d. Full chromophore conjugation with the azide left intact will generate a species less stable than the preceding step and has a predicted absorbance maximum at 451 nm (Supporting Figure S6). The alternative route appears more likely: reduction to the phenylamine followed by loss of the hydroperoxyl group (generating H_2_O_2_). The phenylamine version of hydroperoxyl intermediate has a predicted absorbance maximum of 367 nm. Based on the experimental time course observed in the Figure 1d, the initial species mix dominated by the hydroperoxyl intermediate (predicted λ_max_ = 360 nm) within 1 min of irradiation converts to the dominant hydroperoxyl phenylamine (predicted λ_max_ = 367 nm) that then forms the mature phenylamine chromophore.

### Oxygen access to the chromophore and the twisted Gly67

The structure of im-Venus^66azF^ revealed the presence of O_2_ within the core of the protein close to the I ring of the chromophore. There are two potential ways in which O_2_ becomes located in such a position: during the folding process or diffusion into the core after folding and the cyclisation event. Simulations, whereby O_2_ is replaced by a water molecule, reveal that the twisted conformation of the immature chromophore is more stable by 18.1 kcal/mol than the canonical form (Figure 6a). The presence of water is unsurprising given that O_2_ is not required for the preceding cyclisation and dehydration events. Thus, water helps stabilise the twisted form of Gly67 present in im-Venus^66azF^. If this is the case, O_2_ is likely to displace the water molecule as part of the maturation mechanism, meaning it has to gain access to the core of the protein. Analysis of the im-Venus^66azF^ structure reveals that the “open” His148 conformation may play a role (Figure 6b-c). A channel leads to the chromophore in im-Venus^66azF^ that is only available if His148 occupies the “open” state; when His148 occupies the “closed” state normally observed in mature fluorescent proteins, the tunnel becomes blocked.

**Figure 6.**
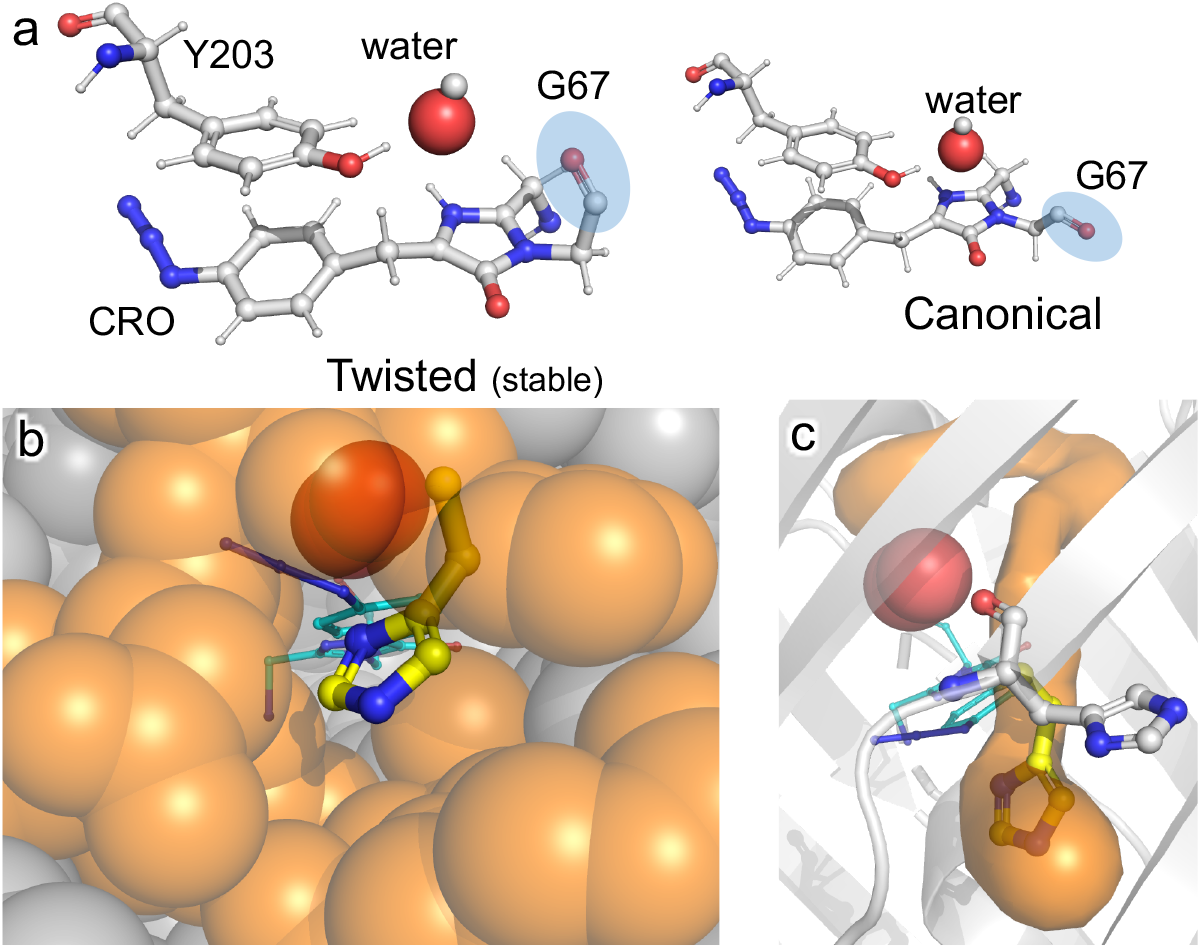
Role of water and internal tunnels in chromophore maturation. (a) Simulation of the twisted conformation stabilised by a water molecule in place of O_2_ (and for reference, the canonical form usually observed in mature Venus^WT^ conformation). The twisted form is more stable by 18.1 kcal/mol compared to the canonical form. The water molecule of interest is highlighted. Chromophore accessibility, as shown by (b) spheres and (c) CAVER tunnel analysis ^37^. The alternative conformation for His148 (yellow sticks) in Venus^WT^ is shown.

## Discussion

Photo-controllable FPs have become an important tool in modern super-resolution cell imaging ^38^. The use of phenyl azide chemistry to control fluorescence, here and in green ^29,30^ and red ^31^ fluorescent proteins, provides a simple and general common mechanism to implement photocontrol across a broad range of the FP colour palette. Indeed, the use of a ncAA in conjunction with a reprogrammed amber stop codon allows secondary control: production of the fusion protein constructs with and without the FP adjunct ^39^. While the demonstration of the conversion of an essentially colourless immature fluorescent protein to an active form through phenylazide photo-decaging confirms the approach as a means to photo-control FPs, insights into the chromophore maturation process are arguably the most important aspect of this work. The one facet our data does confirm is that dehydration occurs before oxidation (Figure 2 and S3), the order of which has been debated in the past ^4^. Based on our work, we propose a maturation route in Figure 7.

**Figure 7.**
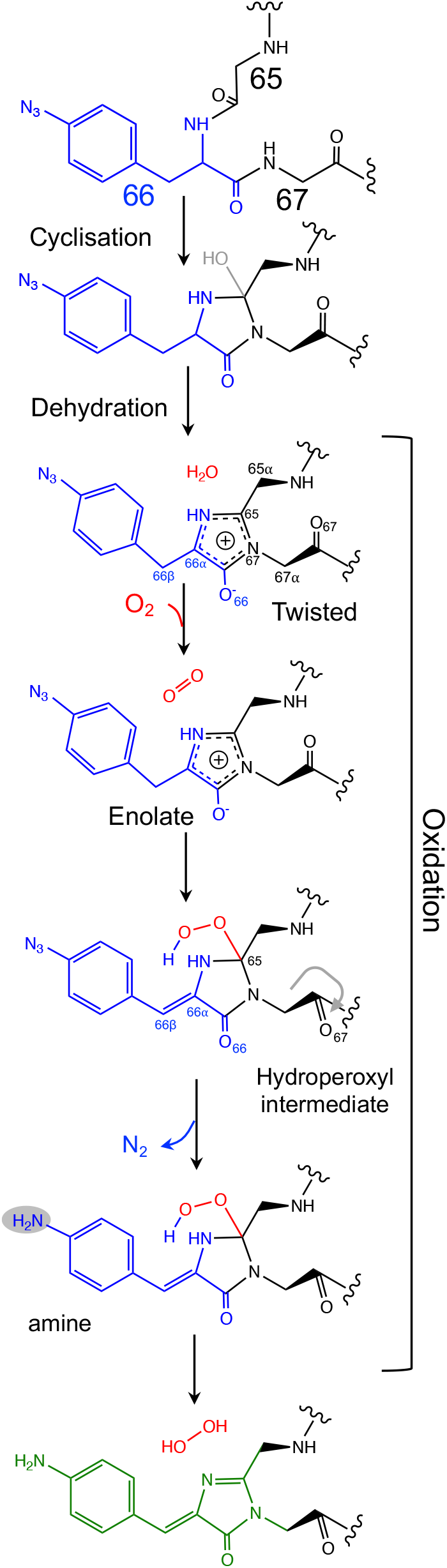
Proposed mechanism for the chromophore maturation, including details of the final oxidation step.

The crystal structure of the im-Venus^66azF^ provides evidence for the nature of an intermediate prior to the final oxidation step: the I ring in the enolate form with the C_66α_ and C_66β_ forming a single bond (Figure 3a). The long C_66_-O_66_ bond is indicative of the enolate whose negative charge is stabilised by critical chromophore maturation residue, Arg96 (Figure 2a) ^14,18,40^. The short C_66_-N_67_ bond (Figure 3a) indicates a C=N bond that would put a positive charge on the nitrogen but the overall bond lengths of the I ring mean a more delocalised system. We don’t preclude a positive charge on the N_66_; our structure indicates there would be no charge clash with Arg96 as suggested previously ^16^.

The crystal structure also provides evidence of the location of O_2_. Two relative positions with respect to the chromophore plane have been proposed: on the Arg96 ^16^ or the Glu222 face ^17^. Here we show that molecular oxygen is placed on the Glu222 face, directly above the I ring (Figure 2). Arg96 has been suggested as the oxygen activator through the positively charged side-chain ^16^ but this unlikely to be the case and may instead play a role in stabilising the enolate form of the I ring.

The twisted Gly67 configuration is clearly observed in the crystal structure and differs from the canonical position normally present in FPs (Figure 2). Glycine has a less restricted ψ angle range. In Venus, the Gly67 ψ dihedral angle is −23° compared to 167° in im-Venus^66azF^, with twisted conformation energetically less favourable (by 7 kcal/mol) when O_2_ is present and also hinders access of oxygen to the I ring (Figure 2). The twisted conformation could be a legacy of the cyclisation reaction whereby the nucleophilic attack of N_67_ on C_65_ will require rotation of the Gly67 ψ angle leading to its observed placement in im-Venus^66azF^. The configuration could be stabilised by the O_67_ H-bonding with the C_65_ hydroxyl group (observed by Getzoff and colleagues ^15^) before condensation. The simulation data reveal that water can stabilise the twisted conformation over the canonical form (Figure 5a), thus suggesting that a hydrated form precedes the oxygen bound step. It is interesting to speculate that the origin of the water molecule may be the product of the cyclisation/dehydration reaction that precedes oxidation. Indeed, while our crystal structure of im-Venus^66azF^ strongly indicates O_2_ is present, we cannot rule out that a minority population of the structures has a water molecule present in the same position.

If a water molecule originally stabilised the twisted Gly67 conformation, this suggests that O_2_ needs to access the protein core for the final oxidation step to take place. We suggest that His148 plays a key role in this process. His148 is dynamic ^41^ and has been observed in both the “open” and “closed” conformations with the former not normally reported in the crystal structure as it is a minor component when observed (for example Arpino *et al* ^42^, Reddington *et al* ^30^ and Brejc *et al* ^43^); the closed conformation is the major form observed in mature FPs as His148 in this configuration H-bonds to the chromophore and plays a critical role in function ^1,9,44,45^. Im-Venus^66azF^ almost exclusively exists in the open conformation (Figure 2a and S3) that generates a channel through to the chromophore (Figure 6b-c). Such a tunnel at a similar position has been observed previously for GFP-like proteins ^46,47^. Thus, His148 may acts as a “gatekeeper” residue, so determining access to the chromophore as well as its functional role ^41^. Given that oxidation is the rate-limiting step in maturation ^4^, it is interesting to speculate that the exchange rate between the two His148 conformations may play a role in defining this rate.

Simulations predict that the next dominant form is hydroperoxyl intermediate attached to C_65_ and not C_66α_, as suggested by others ^16^. The argument against attachment to C_66α_ as the intermediate comes from the observed spectra data (Figure 2b), whereby the dominant 350-360 nm peaks for the intermediate suggests some extension of the conjugated double bond system (here proposed to be from the phenyl azide to C_66α_-C_66β_; Figure 5). The formation of a hydroperoxyl intermediate at C_66α_ would prevent the formation of the double bond with C_66β_. The electron lone pair on the C_65_ hydroperoxyl moiety also aligns perfectly with the two C-N bonds that, in turn, helps stabilise the five-membered I ring. We propose that the nearby Glu222 plays a vital role acting as a general acid/base during the formation of the hydroperoxyl intermediate through the first abstraction and then the donation of a proton to N_66_ (Figure S5). The importance of Glu222 to maturation has been observed previously, with the E222Q mutation in EGFP considerably slowing maturation ^18,40^. During this process a peroxyanion is formed, which abstracts the proton from the activated C_66β_ to form hydroperoxyl intermediate (Figure 4 and S5). Thus, the formation of the C_66α_=C_66β_ double bond occurs before the generation of H_2_O_2_ and not concurrently (Figure 7).

The final and rate-limiting step in the process is the formation of the fully conjugated fluorescent chromophore. In our system, we believe this is a UV induced phase that happens in two steps due to the presence of the azido group: 1. Conversion of the phenyl azide to the phenylamine; 2. Loss of the hydroperoxyl group so generating a fully conjugated system. In our proposed model, we suggest that the reduction of the azide to an amine occurs first (Figure 7). This is based on the predicted absorbance of each species (Figure S6a) and on the expectation that azide conversion to strongly donating amine group would significantly help with loss of the hydroperoxyl moiety (see Figure S6b for mechanistic details). Furthermore, the departure of the OOH group needed for the conversion of hydroperoxyl intermediate to the final maturated chromophore is expected to be greatly facilitated when azide, a mild acceptor (Hammett parameter σ_p_=+0.08), is changed to NH_2_, a strong donor (σ_p_=-0.66). In the azide-substituted peroxide, the lone pair of N_66_ is not able to fully assist in the departure of the OOH group as its electron density is partially delocalized in the other direction, towards the azide. This stereo-electronic tug-of-war is removed once the amine is formed. As the NH_2_ group is a powerful donor, the electron density is no longer shifted from N_66_ to the aryl ring; the lone pair on N_66_ is now free to stabilize the transition state for heterolytic C…OOH bond scission. A fully conjugated chromophore with the azide group attached has a predicted λ_max_ of 451 nm, whereas the amine version of hydroperoxyl intermediate is 367 nm. Given the observed absorbance time course in figure 2 goes from a mixed species with two peaks between 340-360 nm to a single species at 360 nm that directly converts to the single species at 440 nm, the logical progression of the predicted spectra are 360 nm (phenyl azide/hydroperoxyl form) to 367 nm (phenylamine/hydroperoxyl forms) to 444 nm (mature amine chromophore).

## Conclusion

For the first time, our X-ray structure of im-Venus^66azF^ provides an experimental insight into the role played by oxygen in fluorescent protein chromophore maturation. The fortuitous trapping of oxygen between Tyr203 and an alternative conformation of the strictly conserved Gly67 before photo-decaging of azF66 was critical. From our structure and subsequent modelling, the cyclisation-dehydration-oxidation model mechanism is the likely route to chromophore maturation but with new insights concerning the nature of the enolate intermediate together with mechanistic details of the O_2_-dependent steps. These new insights not only improve our fundamental understanding of chromophore maturation but may help aid our ability to generate improved fluorescent protein for future applications.

## Methods and materials

### Engineering, production and structure of Venus^66azF^

The generation of the mutant and subsequent recombinant production of Venus^66azF^ is outlined in the supporting information and based on a previously published procedure ^47^. Protein production and purification was carried out in the dark to prevent photolysis of azF. Crystallisation, diffraction data collection and structure determination of im-Venus^66azF^ was performed as outlined in the supporting information. A long, spiny crystal of im-Venus^66azF^ was observed after a few days in the A12 condition of the PACT premierTM HT-96 screen (0.01 M Zinc chloride, 0.1 M Sodium acetate, pH 5 and 20% w/v PEG 6000). The structural statistics are provided in Supporting Table 2.

### Absorbance and fluorescence spectroscopy

Absorbance spectra were recorded using a Cary 60 spectrophotometer (Agilent) in a 1 cm pathlength quartz cuvette. The molar absorbance coefficients were calculated by recording absorbance spectra of Venus samples with a known concentration (5-10 μM) and then extrapolated to 1 M using the Beer-Lambert law. Emission spectra were recorded using a Cary Eclipse fluorimeter (Varian) using a 5 mm x 5 mm QS quartz cuvette. Protein concentration of 0.5 μM for Venus^WT^ or 10 μM for Venus^66azF^ was used with a fixed scan rate of 120 nm/min with a 5 nm slit width. The excitation wavelength for Venus^WT^ and Venus^66azF^ were determined from their absorbance spectrum. Bacterial live cell imaging was performed as outlined in the Supporting Information.

### Photolysis of Venus^66azF^

Photolysis experiments were carried out using a UVM-57 Handheld UV lamp (6 W; 302 nm UV, UVP Cambridge, UK) and 1 cm pathlength quartz cuvette (Hellma), essentially as described previously ^29,46^. A maximum of 500 μl of protein sample (10 μM) was pipetted into a cuvette and exposed to the UV (302 nm) for the indicated periods of time at a distance of 1cm. The absorbance spectra and emission spectra were recorded immediately afterwards, as described above.

### QM/MM Simulations

Coordinates of heavy atoms of the im-Venus^66azF-^ crystal structure were used to construct a full atom three-dimensional model system. Previous examples in quantum mechanics/molecular mechanics (QM/MM) simulations of the mechanism of chromophore maturation included the wild-type GFP^17^, its Gly65-Gly66-Gly67 mutant ^48^, and the recent modeling of Dreiklang ^49^, a photoswitchable protein with the chromophore formed from the same amino acid residues Gly65, Tyr66, Gly67, as in Venus. Hydrogen atoms were added manually using molecular mechanics tools; the side chains of Arg and Lys were assumed as positively charged, the side chains of Glu and Asp as negatively charged. The model protein molecule was fully surrounded by explicit water molecules.

Structures of possible intermediates in the maturation reaction were optimized in QM/MM calculations. A large fraction of the chromophore-containing pocket was assigned to the QM-part. The Gly65-azF66-Gly67 fragment of the immature chromophore (CRO), the side chains from Arg96, Tyr203, Ser205, Glu222 and 4 water molecules were included. This initial composition was considered to model structures without the oxygen molecule. In majority of calculations, the O_2_ species was inserted to the cavity near CRO. Calculations of energies and energy gradients in QM were carried out using Kohn-Sham DFT with the PBE0 functional ^50^ and the cc-pVDZ basis set. The AMBER force field was used in MM. The NWChem software package ^51^ was applied to scan fragments of potential energy surface. To model the system in the triplet electronic state, the unrestricted DFT approach was used. Vertical excitation energies at selected points on the ground state potential energy surface were computed using the extended multiconfigurational quasi-degenerate perturbation theory in the second order (XMCQDPT2) ^52^ the protocol that we verified earlier and used extensively in studies of the photoreceptor proteins ^53^. Here, the perturbation theory calculations were based on the complete active space self-consistent field (CASSCF) wavefunctions obtained by distributing 16 electrons over 12 orbitals and using density averaging over 15 states. To perform these calculations using the Firefly quantum chemistry package ^54^, large molecular clusters including the QM parts of the system were selected. Natural Bond Orbital (NBO) analysis was used to evaluate stereoelectronic interactions ^55,56^. Geometry optimizations for NBO evaluations were performed with (SMD ^57^=water) for solvation corrections and the unrestricted *w*B97X DFT functional ^58^ (with an integration grid of pruned 175,974 for first-row atoms and 250,974 for atoms in the second and later rows) with the 6-311 ++G(2d,p) basis set for all atoms. Grimme’s D2 version for empirical dispersion ^59,60^ was also included. Natural Bond Orbital (NBO) analyses were performed with *NBO6* linked to *Gaussian 16*. They were used to gauge the magnitude of the hyperconjugative interactions in the presented systems.

## Acknowledgements

We would like to thank the staff at the Diamond Light Source (Harwell, UK) for the supply of facilities and beam time, especially Beamline I03 and I04 staff, under beamtime code mx18812. We thank BBSRC (BB/H003746/1 and BB/M000249/1), EPSRC (EP/J015318/1). H.S.A. was supported by the Higher Committee for Education Development in Iraq. We would like to thank the Protein Technology Hub, School of Biosciences, Cardiff University for use of facilities. B.L.G. and I.V.P. thank the Russian Science Foundation (project 17-13-01051) for support of the modelling part of this work. The calculations were carried out using the equipment of the shared research facilities of HPC computing resources at Lomonosov Moscow State University. The use of supercomputer resources of the Joint Supercomputer Center of the Russian Academy of Sciences is also acknowledged.

## Author contributions

All authors contributed to the writing of the paper and analysing data. HSA. undertook structural biology and functional characterisation the Venus proteins; TH helped prepare mutant proteins. PJR collected structural data and helped with structure determination and refinement. BLG, IVP and AVN. developed computational models and carried out computer simulations. IA and GPG provided analysis of stereoelectronic factors involved in the formation and reactivity of hydroperoxide intermediates. CB analysed data. DDJ conceived and directed the project, and analysed data.

---

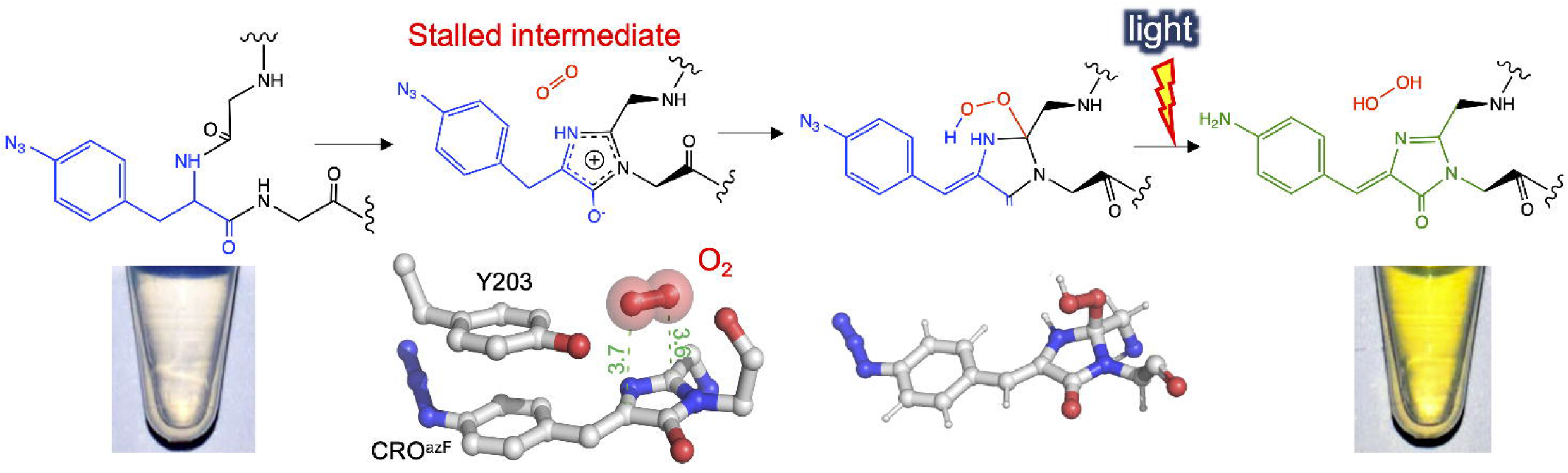

