## Supplementary material for "Stalling chromophore maturation of the fluorescent protein Venus reveals the molecular basis of the final oxidation step": SI methods, tables and figures

### Supporting Methods.

#### Venus$^{66azF}$ gene sequence

ATGCGGGGTTCTCATCATCATCATCATCATGGTATGGCTAGCATGACTGGTGGACAGCAAATGGGTCGGGATCTG
TACGAGAACCTGTACTTCCAGGGCTCGAGCATGGTGAGCAAGGGCGAGGAGCTGTTCACCGGGGTGGTGCCCATC
CTGGTCGAGCTGGACGGCGACGTAAACGGCCACAAGTTCAGCGTGTCCGGCGAGGGCGAGGGCGATGCCACCTAC
GGCAAGCTGACCCTGAAGCTGATCTGCACCACCGGCAAGCTGCCCGTGCCCTGGCCCACCCTCGTGACCACCCTG
GGC**TAG**GGCCTGCAGTGCTTCGCCCGCTACCCCGACCACATGAAGCAGCACGACTTCTTCAAGTCCGCCATGCCC
GAAGGCTACGTCCAGGAGCGCACCATCTTCTTCAAGGACGACGGCAACTACAAGACCCGCGCCGAGGTGAAGTTC
GAGGGCGACACCCTGGTGAACCGCATCGAGCTGAAGGGCATCGACTTCAAGGAGGACGGCAACATCCTGGGGCAC
AAGCTGGAGTACAACTACAACAGCCACAACGTCTATATCACCGCCGACAAGCAGAAGAACGGCATCAAGGCCAAC

TTCAAGATCCGCCACAACATCGAGGACGGCGGCGTGCAGCTCGCCGACCACTACCAGCAGAACACCCCCATCGGC
GACGGCCCCGTGCTGCTGCCCGACAACCACTACCTGAGCTACCAGTCCGCCCTGAGCAAAGACCCCAACGAGAAG
CGCGATCACATGGTCCTGCTGGAGTTCGTGACCGCCGCCGGGATCACTCTCGGCATGGACGAGCTGTACAAGTAA

**Engineering and production of Venus$^{66azF}$**

The Venus plasmid was originally obtained from Addgene (Venus-pBAD, #54859). The introduction of TAG substitution in place of the codon for Tyr66 was performed using whole plasmid (or inverse) PCR with mutagenic primer Venus-TAG-66-F (5'-GGC**TAG**GGCCTGCAGTGCT-3'; the letter in bold represent the introduced TAG mutation) and Venus-TAG-66-R (5'-CAGGGTGGTCACGAGGGTG-3'). Whole plasmid PCR was performed using the gene encoding Venus resident within the pBAD plasmid as template and Q5 DNA polymerase as described previously [1]. Introduction of the mutation was confirmed by DNA sequencing. The recombinant production of azF-containing Venus variants in *E. coli* was performed as described previously [1]. The plasmid pDule-azF was (a kind gift for Ryan Mehl [2] and supplied by Addgene) was used to facilitate incorporation of exogenous azF (0.5 mM) supplemented to 2xYT culture medium. From this point onwards, all stages were performed in the dark or samples wrapped in foil to prevent premature photolysis. Production of Venus was induced by the addition of 0.2% (w/v) arabinose when the culture reached $OD_{600}$ of 0.7. The cells were incubated a further 20 hr at 25ºC. Cells were pelleted and then resuspended in 50 mM Tris-HCl, pH 8 followed by lysis using a French pressure cell press. Venus was purified initially by nickel affinity chromatography using a gravity flow Protino$^{R}$ Ni-TED 2000 affinity column. Venus was eluted using 50 mM Tris-HCl, 250 mM imidazole, pH 8. Pure protein fractions were pooled and polished by size exclusion chromatography using a Hiload$^{TM}$ 16/600 Superdex$^{TM}$ S75 column equilibrated with 50 mM Tris-HCl, pH 8.

**Structure determination of Venus$^{66azF}$**

For dark state Venus$^{azF66}$ sample, a concentrated purified sample (~20 mg/ml) was used for screening crystal formation using the PACT *premier*$^{TM}$ HT-96 screen and JCSG-plus$^{TM}$ HT-96 screen (Molecular Dimensions, UK). The sitting drop vapour diffusion method was used in a sealed tray and kept at 20°C. All the steps of crystal preparation were performed in the dark. The growth of crystals was monitored under a light microscope, and grown crystals were harvested by picking individual crystals in a mounted litholoop (Molecular Dimensions) and plunging them into liquid nitrogen. Diffraction data were collected at the Diamond Light Source, Harwell, UK, at beamline I04. Data were reduced with the XIA2 package [3], POINTLESS was used for space group assignment, scaling and merging were completed with Aimless [4] and TRUNCATE [5]. Molecular replacement with PHASER was used to solve the structure of Venus$^{66azF}$ dark state using sfGFP$^{66azF}$ dark state (PDB accession 4J88[6]) as a model[7]. The COOT program was used to adjust the structure manually for several repeated cycles, TLS restrained refinement was used to refine the structure using RefMac [8]. The CCP4 package was used for the above routines [5].

**Bacterial live cell imaging.**

Widefield fluorescence microscopy was used for bacterial live cell imaging. For photoactivation of Venus$^{Y66azF}$, *E. coli* Top10 cells were first induced to express Venus$^{66azF}$. A Coverwell$^{TM}$ imaging chamber (Sigma-Aldrich) was fixed on a glass slide and 0.2 ml of 1% agarose was applied before the induced bacterial cells (0.2 ml) were added and covered with a cover slide. A sample of uninduced bacterial cells was prepared in the same manner as a control. Slides for induced and uninduced bacterial cells were exposed to UV-light for 1 min at a distance of about 1 cm. Transmitted and widefield fluorescence images visualised using an inverted Olympus IX73 widefield fluorescence microscope. Images were collected with a Hamamatsu Orca flash 4.0 camera with x100 objective lens using HCImaging software and a Prior Lumen200Pro light source. Fluorescence emission was separated by a multiband dichroic emission filter set 69002 (Chroma), a wavelength of 450 nm was used for the excitation.

**Supporting Table S1. Spectral characteristics of fluorescent proteins.**

| Variant | $\lambda_{max}$ (nm) | | $\lambda_{em}$ (nm) | | $\varepsilon$ ($M^{-1}cm^{-1}$) | | QY | |
|---|---|---|---|---|---|---|---|---|
| | Dark | UV | Dark | UV | Dark | UV | Dark | UV |
| [a] Venus$^{WT}$ | 515 | N/A | 528 | N/A | 92200 | N/A | 0.45 | N/A |
| Venus$^{66azF}$ | 352 | 440 | N/A | 511 | 30260 | 18500 | N/A | 0.1 |
| [b] sfGFP$^{66azF}$ | 391 | 440[c] | 500 | 500 | 42870 | 24620 | 0.13 d | 0.16 |

[a] data from [1]; [b] data from [6]; [c] final chromophore maturation product is the phenyl amine; [d] excitation at 446 nm, where $\varepsilon$ is 2300 $M^{-1}cm^{-1}$

**Supporting Table S2.** Crystal structure diffraction and refined statistics of dark state Venus$^{66azF}$

| PDB Code | 6SM0 |
|---|---|
| **Data collection/reduction statistics** | |
| Diamond Beamline | I04-1 |
| Wavelength | 0.91587 |
| *a, b*, *c* (Å) | 50.947, 63.251, 69.915 |
| Space group | P $2_1$ $2_1$ $2_1$ |
| Resolution (Å) | 1.909 – 63.26 |
| Outer shell | 1.909 – 1.96 |
| *R*-merge (%) | 16.4 (114.5) |
| CC1/2 | 0.989 (0.355) |
| I / σ | 7.4 (1.8) |
| Completeness (%) | 99.9 (99.9) |
| Total Measurements | 103,864 (4,920) |
| Unique Reflections | 18,146 (1,317) |
| Wilson B-factor(Å$^2$) | 16.0 |
| **Refinement Statistics** | |
| Non-H Atoms | 2,118 |
| R-work reflections | 17,175 |
| R-free reflections | 1,238 |
| R-work/R-free | 19.4 / 24.2 |
| **rms deviations** | |
| Bond lengths (Å) | 0.013 |
| Bond Angles (°) | 1.854 |
| [1]Coordinate error | 0.135 |
| Mean B value (Å2) | 19.6 |
| **Ramachandran Statistics** | |
| Favoured/allowed/Outliers | 187 / 5 / 0 |
| % | 97.4 / 2.6 / 0.0 |

**Supporting Table 3.** Chromophore bond distances.

| Bond | Venus | im-Venus$^{66azF}$ |
|---|---|---|
| Imidazolone (I) ring (in Å) | | |
| $C_{66}$-$O_{66}$ | 1.20 | 1.48 |
| $C_{66}$-$C_{66\alpha}$ | 1.49 | 1.42 |
| $C_{66\alpha}$-$N_{66}$ | 1.43 | 1.34 |
| $N_{66}$-$C_{65}$ | 1.41 | 1.33 |
| $C_{65}$-$N_{67}$ | 1.36 | 1.34 |
| $N_{67}$-$C_{66}$ | 1.36 | 1.26 |
| $N_{67}$-$C_{67\alpha}$ | 1.48 | 1.44 |
| $C_{65}$-$C_{65\alpha}$ | 1.50 | 1.48 |
| Methylene bridge P ring (in Å) | | |
| $C_{66\alpha}$-$C_{66\beta}$ | 1.35 | 1.45 |
| $C_{66\beta}$-$C_{66\gamma}$ | 1.49 | 1.47 |
| Additional bonds (in Å) | | |
| $C_{67}$-$O_{67}$ | 1.22 | 1.36 |
| $C_{67\alpha}$-$C_{67}$ | 1.48 | 1.50 |

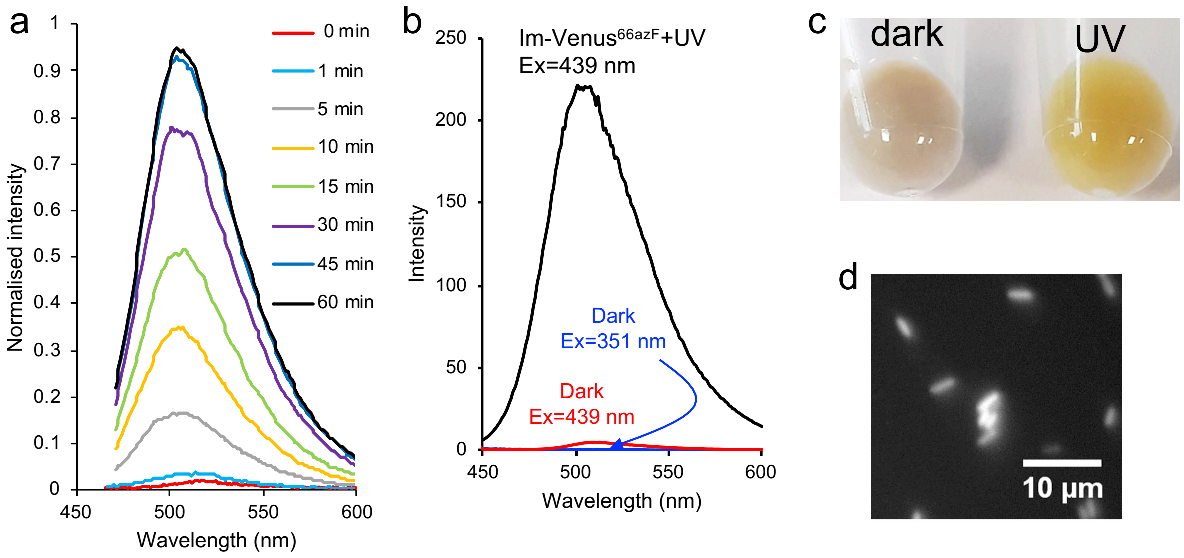


**Supporting Figure S1.** (a) Time course of Venus$^{66azF}$ fluorescence emission (on excitation at 435 nm) on irradiation for different time points indicated in the figure. (b) Emission profile of im-Venus$^{66azF}$ on excitation at 351 nm (blue) and 439 nm (red) before UV exposure and emission on excitation at 439 nm after UV exposure (black). (c and d) *in situ* activation of Venus$^{66azF}$ with *E. coli* cell cultures before and after UV irradiation (c) and wide-field imaging of *E. coli* cells after activation with UV (d).

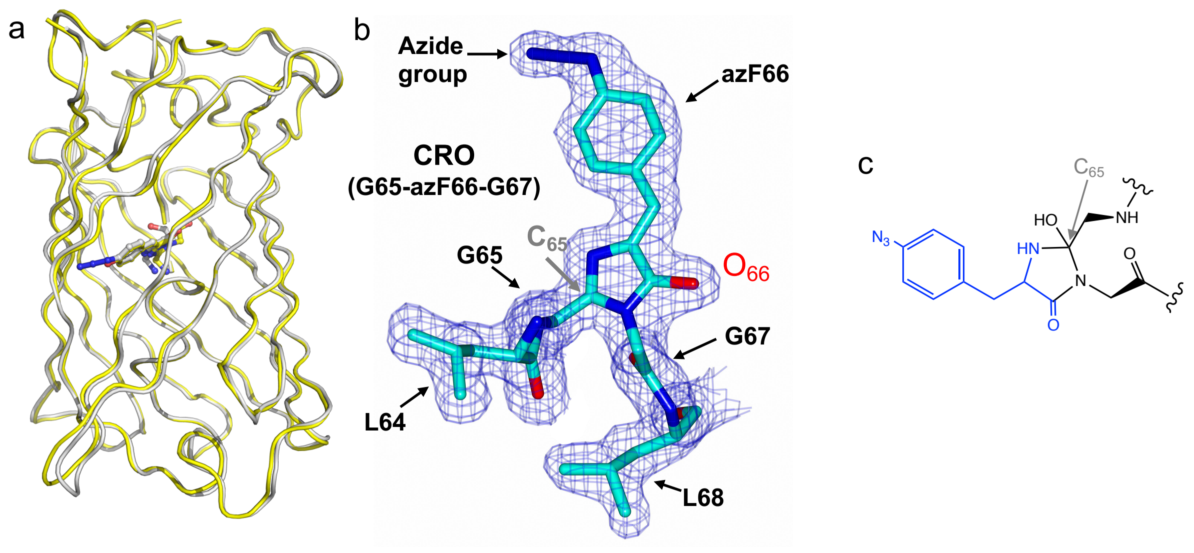


**Supporting Figure S2.** (a) Overlay of the Venus$^{WT}$ (yellow) and im-Venus$^{66azF}$ (grey). The chromophore is shown as sticks. The RMSD between backbone is 0.39 Å. Chromophore structure of im-Venus$^{66azF}$. (b) Electron density map (2Fo-Fc, 1.5 sigma) of the CRO. Each residue component of the original sequence is shown. The $C_{65}$ carbon where a hydroxyl group would be observed prior to the dehydration step is outlined; no electron density for the expected hydroxyl group is observed. The elongated $C_{66}$-$O_{66}$ bond is indicated by the $O_{66}$ annotation. (c) Chemical structures of the chromophore before the hydration step.

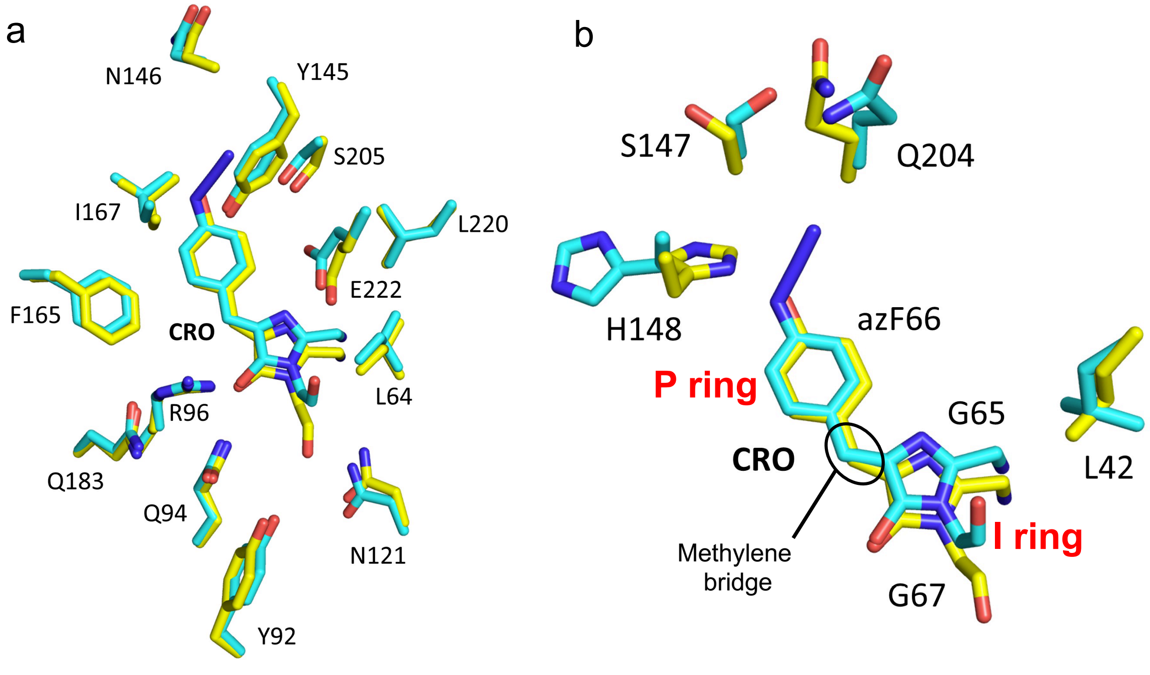

**Supporting Figure S3.** Comparison of residue positions between Venus$^{WT}$ (yellow) and im-Venus$^{66azF}$ (cyan). Panels a and b show different residue sets for clarity, with residues undergoing a larger conformational change highlighted in b.

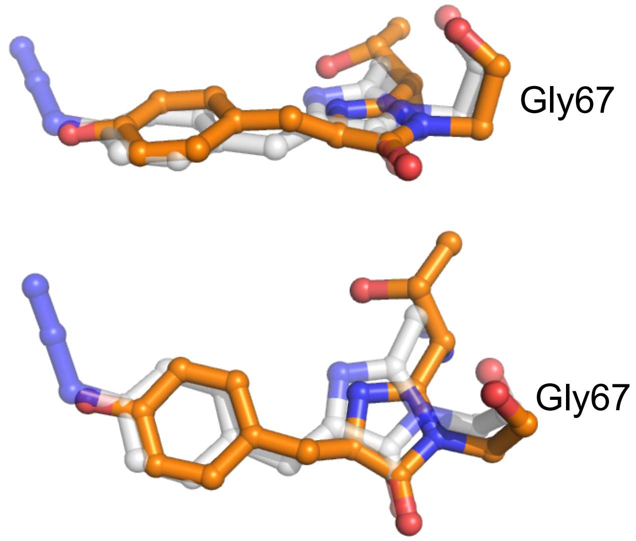


**Supporting Figure S4.** Comparison of the chromophore structure of im-Venus$^{66azF}$ (grey) and unpublished GFP mutant Q183E (PDB 2qt2; orange). The GFP mutant has a partially formed chromophore and the same orientation of the Gly67 carbonyl oxygen observed for im-Venus$^{66azF}$.

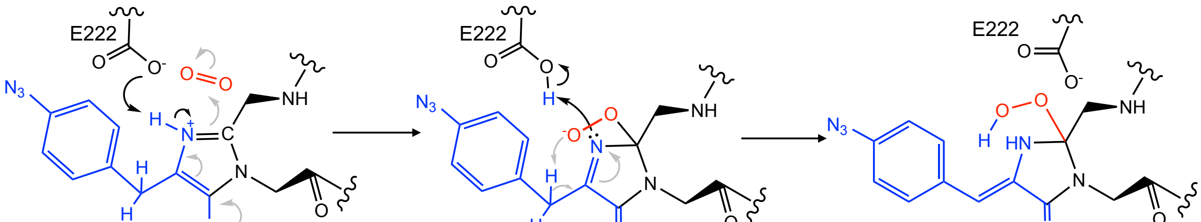


**Figure S5.** Schematic outline of the reaction of $O_2$ with im-Venus$^{66azF}$, including the potential intermediates and the role of residue E222.

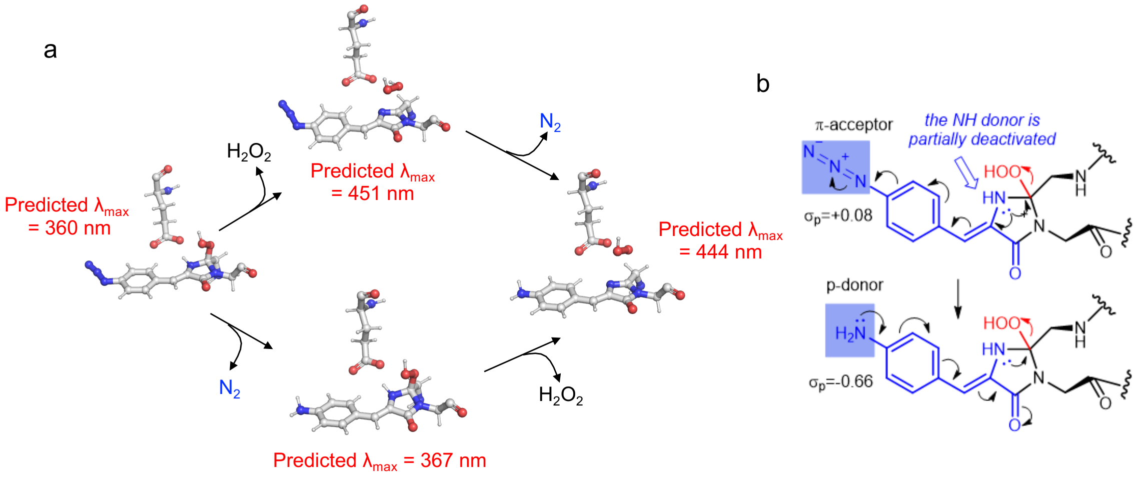


**Supporting Figure S6. (a)** Potential routes to mature phenyl amine chromophore from the hydroperoxyl intermediate. The two routes are chromophore conjugation followed by azide reduction and *vice versa*. The predicted $\lambda_{max}$ values are shown. It should be pointed out that the presumed species without the hydroperoxyl and phenyl azide present ($\lambda_{max}$=451 nm) is considerably higher in energy than the hydroperoxyl intermediate. Therefore, the upper route is unlikely, as also clarified in the panel (b). **(b)** Mechanistic analysis highlights the contrasting effects of azide and amine substituents on the elimination of hydroperoxide in the final aromarisation step. The decelerating effect of the acceptor azide on the OOH elimination is deactivated once the azide is converted into a strongly donating amine moiety
